# Analysis of the genetic basis of height in large Jewish nuclear families

**DOI:** 10.1101/303008

**Authors:** Danny Zeevi, Joshua S. Bloom, Meru J. Sadhu, Adi Ben Yehuda, David Zangen, Ephra Levy-Lahad, Leonid Kruglyak

## Abstract

Despite intensive study, most genetic factors that contribute to variation in human height remain undiscovered. We conducted a family-based linkage study of height in a unique cohort of very large nuclear families from a founder (Jewish) population. This design allowed for increased power to detect linkage, compared to previous family-based studies. We identified loci that together explain an estimated 6% of the variance in height. We showed that these loci are not tagging known common variants associated with height. Rather, we suggest that the observed signals arise from variants with large effects that are rare globally but elevated in frequency in the Jewish population.

## Introduction

Height is a classic genetically complex quantitative trait with high heritability (~80%-90%^1,2^). Despite intensive study, the genetic basis of variation in height remains mostly unexplained. Genome-wide association studies (GWAS) in hundreds of thousands of individuals have identified hundreds of common variants significantly associated with height ^3,4^. However, the individual effect sizes of these variants are small, and all variants identified to date jointly explain only ~20% of the heritability of height. One proposed explanation for the gap between the overall heritability of height and that explained by common variants is the contribution of rare genetic variants with large phenotypic effects ^5^. Recent studies lend support to this idea by identifying such variants and showing that their effect sizes are inversely proportional to their frequencies ^6,7^ Several examples of large-effect variants that are rare globally but are more common in certain founder populations have been reported for height (in Sardinians ^8^ and in Puerto Ricans ^9^) and diabetes (in Greenlanders ^10^), However, identifying associations between rare variants and traits of interest typically requires very large sample sizes ^11^. For instance, 750,000 participants were required to identify 32 rare variants (those with frequency <1%) that affect height ^7^.

Rare variants can in principle be identified in family-based linkage studies with lower sample size requirements than association studies. 21 family-based studies of height have been conducted, mostly prior to the GWAS era ^12–32^. Only a few of these studies detected Quantitative Trait Loci (QTLs) linked to height at the accepted level of genome-wide statistical significance, with the three most convincing findings all reported by a single study with the largest sample size of subjects from one ethnicity ^23^. Moreover, few if any of the reported QTLs were replicated across multiple studies^14,23,32^.

We sought to overcome some of the limitations of previous linkage studies by focusing on several factors that influence statistical power, including the minor allele frequency (MAF) of causal variants, the strength of linkage between causal and typed variants, quality of genotyping and phenotyping, and the effect size of the variant ^33,34^ Specifically, we attempted to increase power by studying the genetic architecture of height in very large nuclear families (10 to 20 siblings per family) from a founder (Jewish) population. We acquired highly accurate measurements of height, and we used dense genotyping arrays to fully reconstruct the inheritance patterns in these families. The use of large pedigrees drawn from a population with a small effective population size should increase the effective allele frequency of variants of interest, enabling their detection in a study with a modest sample size.

## Results

### Increasing effective allele frequencies by studying very large nuclear families

To increase the power to detect effects of rare variants, we sought to increase their effective frequency in our cohort by studying very large nuclear families. A rare variant carried by a parent of such a family automatically rises to a frequency of ~25% among the children. However, this effect is rapidly diluted when many unrelated small families are combined, as was done in all previous family studies of height. To minimize such dilution, we recruited 397 participants from 29 very large nuclear families containing 10 to 20 siblings per family (mean=12 siblings). In addition, siblings in eight of the nuclear families have first cousins in several other nuclear families in the cohort. Any variant segregating in our cohort has a minimum expected MAF of ~1% if it is present in only one family, and higher if it is present in multiple families, regardless of its frequency in the general population.

### Increasing allele frequencies by studying a founder population

Another approach to increase MAF is to study a founder population, where some variants that are rare in a cosmopolitan population can rise to high frequencies due to a small number of founders and subsequent genetic drift. We therefore recruited our cohort from the Jewish population. Specifically, 80% of our cohort consists of Ashkenazi Jews (AJ). Today ‘s 10 million AJ have an effective population size (N_e_) of ~350 as a result of a founder event ~700 years ago^35^. This effective population size is small even compared to other founder populations such as Finland (N_e_~3000) ^36^ and Iceland (N_e_~5000) ^37^ Two unrelated AJ on average share ~30 times more of their genome in long (>3Mbps) identity by descent (IBD) segments than two unrelated non-Jewish Caucasians ^38^. As a result, variants that are rare in other populations can rise to high frequencies in AJ. Indeed, at least 40 Mendelian genetic diseases in AJ are caused predominantly by such founder mutations that are non-existent or rare in other populations ^39,40^. The other 20% of our sample are Jews of other ethnicities. Previous studies showed that Jews who are not Ashkenazi are on average closer genetically to Ashkenazi Jews than to their non-Jewish neighbors ^41^.

To estimate the shift in allele frequencies between the European and AJ populations, we compared allele frequencies in whole genomes of 7509 non-Finnish Europeans and 151 Ashkenazi Jews, both from the gnomAD database ^42^ (**Fig. 1**). The database contains 90.5 million bi-allelic high quality variants found in Europeans, and 82.5 million (91%) of these are rare (MAF<1%). 95% of the rare variants found in Europeans are not observed in the AJ sample, but 33,331 variants that are rare or not observed in Europeans are common in AJ (allele frequency ≥5%), and 757 of them reach allele frequency ≥10%. To test whether the increase in the frequency of rare variants in AJ is a consequence of small sample size, we used the European allele frequencies as probabilities for randomly sampling 151 Europeans from gnomAD and repeated the analysis. No rare variant reached a frequency of ≥5% in this sub-sample (**Fig. 1b**).

**Fig. 1.**
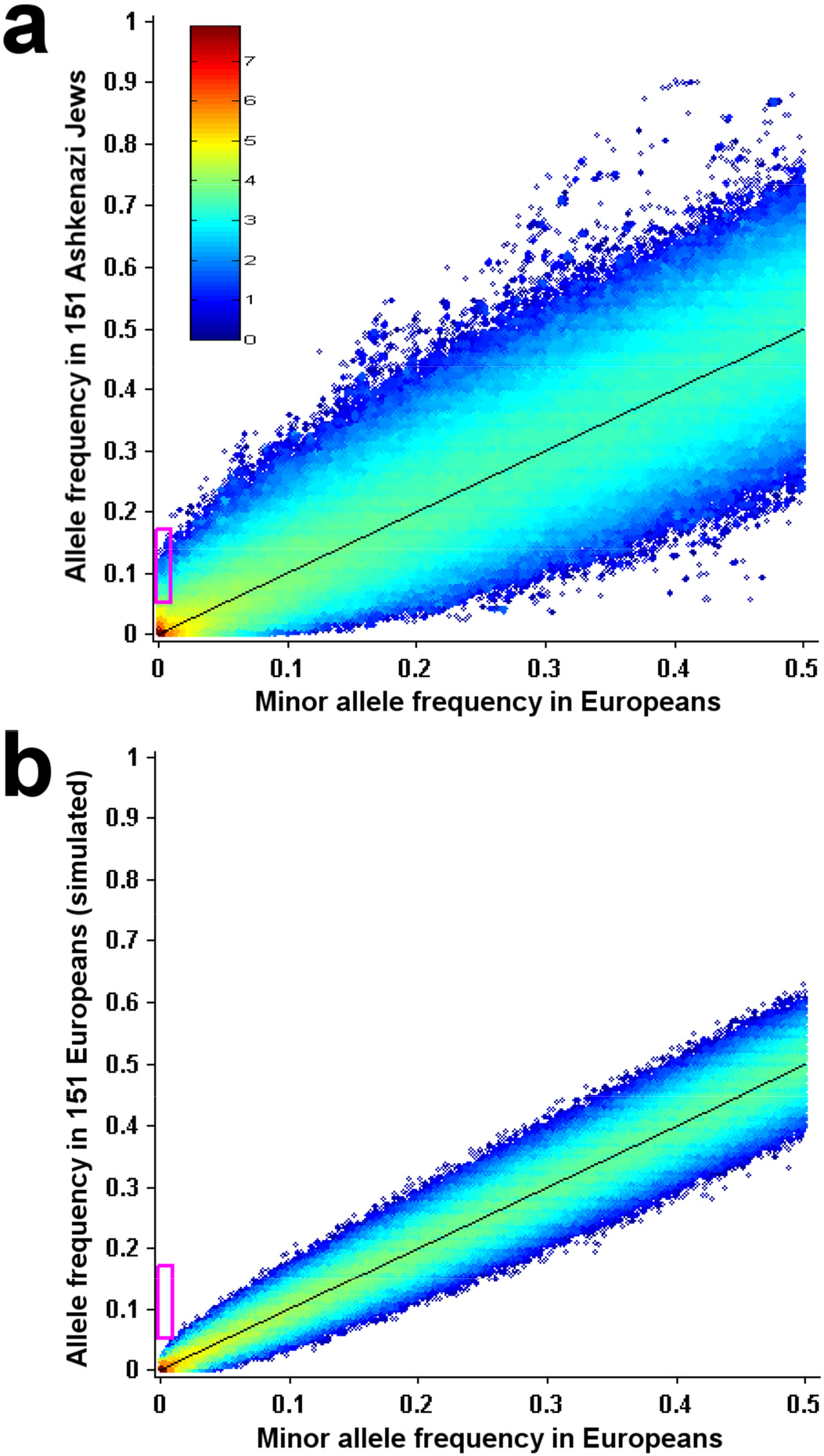
Founder effect in Ashkenazi Jews. **(a)** Minor allele frequency of variants in 7509 Europeans (X-axis) vs. their allele frequency in 151 Ashkenazi Jews (Y-axis). Each dot represents one of ~90.5 million genetic variants from the gnomAD database. Color displays density, i.e. the number of variants in that region of the plot, on a log10 scale. The magenta rectangle encloses all variants that are rare (MAF<1%) in Europeans but common (MAF≥5%) in Ashkenazi Jews. Black line shows y=x. **(b)** Same as in (a), but with allele frequency in a random sub-sample of 151 Europeans shown on the Y-axis.

Increasing the MAF of a variant increases the power to detect that variant in several ways. First, power is dependent directly on MAF (See **Methods M4** for power calculations). Second, rare variants tend to have a lower quality of genotyping and imputation ^33^, and increasing MAF allows for a higher quality of both. Third, the effect sizes of variants are typically inversely proportional to their frequencies in cosmopolitan populations ^6,7^, and therefore causal variants that are rare elsewhere but are common in our study population may have larger effect sizes. The combination of these factors, together with our use of large families and accurate phenotyping (see below), has the potential to increase power to detect Quantitative Trait Loci (QTL) that influence height.

### Phenotype accuracy

Most previous height studies used measurements acquired incidentally as part of studying other traits or diseases. Such measurements may suffer from low accuracy. To ensure phenotype accuracy and increase statistical power, we measured the height of each participant to the nearest 0. 1 cm, with four technical repeats, and with one researcher (D.Ze.) conducting all measurements and using the same measurement system. We tested different methods to correct heights for age and sex (**Methods M1-M3**) and used a non-linear correction for age that maximized heritability in our sample.

### Multiple siblings and dense SNP arrays enable reconstruction of fully informative inheritance patterns for linkage analysis

Nearly all previous family linkage studies of height ^12–32^ used sparse maps of microsatellite markers, which provide incomplete inheritance information content and a corresponding reduction in statistical power to detect QTL ^43^. A dense map of single nucleotide polymorphisms (SNPs), combined with multipoint linkage analysis, can generate near-perfect information content ^43^, but existing computational tools are not well-suited to carry out multipoint analysis with hundreds of thousands of markers. To obtain fully informative inheritance patterns at every position in the genome, we genotyped ~630,000 SNPs and developed a method for identity-by-descent (IBD) reconstruction that leverages information from the large number of siblings in each family (**Methods M6**). Briefly, we compared genotypes of each sibling pair to identify IBD segments, and then used these segments identified across all pairs to reconstruct, at every genomic position, the fully informative inheritance pattern of all four parental haplotypes in the children (**Fig. S3**). Every recombination event in a sibling was identified as a change in his or her IBD relationship pattern with the other siblings. Because the number of recombination events per chromosome per participant (typically 1-2) is much smaller than the number of siblings in each of our families, we were able to identify these IBD switching events with high certainty, even in the absence of parental genotypes. To test the accuracy of the method, we calculated the average pair-wise IBD sharing for the genome. Two siblings shared zero, one or two alleles IBD for 24.9%, 50% and 25.1% of the genome, compared to the theoretical expectation of 25%, 50% and 25%. The IBD segments identified by our method provide near-perfect inheritance information for linkage analysis between genomic loci and height.

### QTL mapping

To identify regions of the genome that co-segregate with height differences (QTLs), we conducted linkage analysis by contrast tests (ref. ^44^, **Methods M7-M9**). Briefly, at every position in the genome, we compared the heights of siblings that inherited one of the two possible haplotypes from a given parent to the heights of those who inherited the other haplotype. We used permutations of height among siblings in a family to calculate significance, which we express as an equivalent LOD score (**Fig. S4**). It has been shown ^45^ that for sib pair analysis, genome-wide significance at an FDR of 5% corresponds to LOD≥3.6 (equivalent to p-value < 2×10^-5^). We identified two significant QTLs, on chr14 (LOD=4.13) and chr17 (LOD=3.8). Permutation analysis showed that 0.11 QTLs are expected by chance at this threshold, which corresponds to an empirical FDR of 5.5%, in close agreement with the theoretical value.

We reasoned that because height is a complex trait that is governed by multiple loci with relatively small effects, an FDR analysis of larger sets of loci with lower locus-specific significance would be appropriate and could increase the sensitivity of our study. This approach is frequently employed in GWAS ^46^, but was not attempted in previous linkage studies of height. To carry out such analysis, we compared the number of observed QTLs for a range of LOD score thresholds below 3.6 to the number expected in the absence of true signals, determined from permutations. As expected, lower thresholds resulted in larger numbers of detected QTLs, with an increasing FDR (**Fig. 2a and 2b**). Importantly, the total number of detected QTLs remained significant (P<0.05 compared to permutations, **Fig. 2c**) from the strictest threshold (LOD=3.6; 2 QTLs; P=0.009; FDR = 5.5%), down to LOD=1.1 (24 QTLs; P=0.019; FDR = 68%; **Supp. Table T1**). We used the FDR to estimate the number of true QTLs as a function of decreasing detection thresholds (**Fig. 2a**). The number of true QTLs maximizes at LOD=1.3, where we estimate 8.8 true QTLs out of 20 that were detected (P=0.002; FDR=56%).

**Fig. 2.**
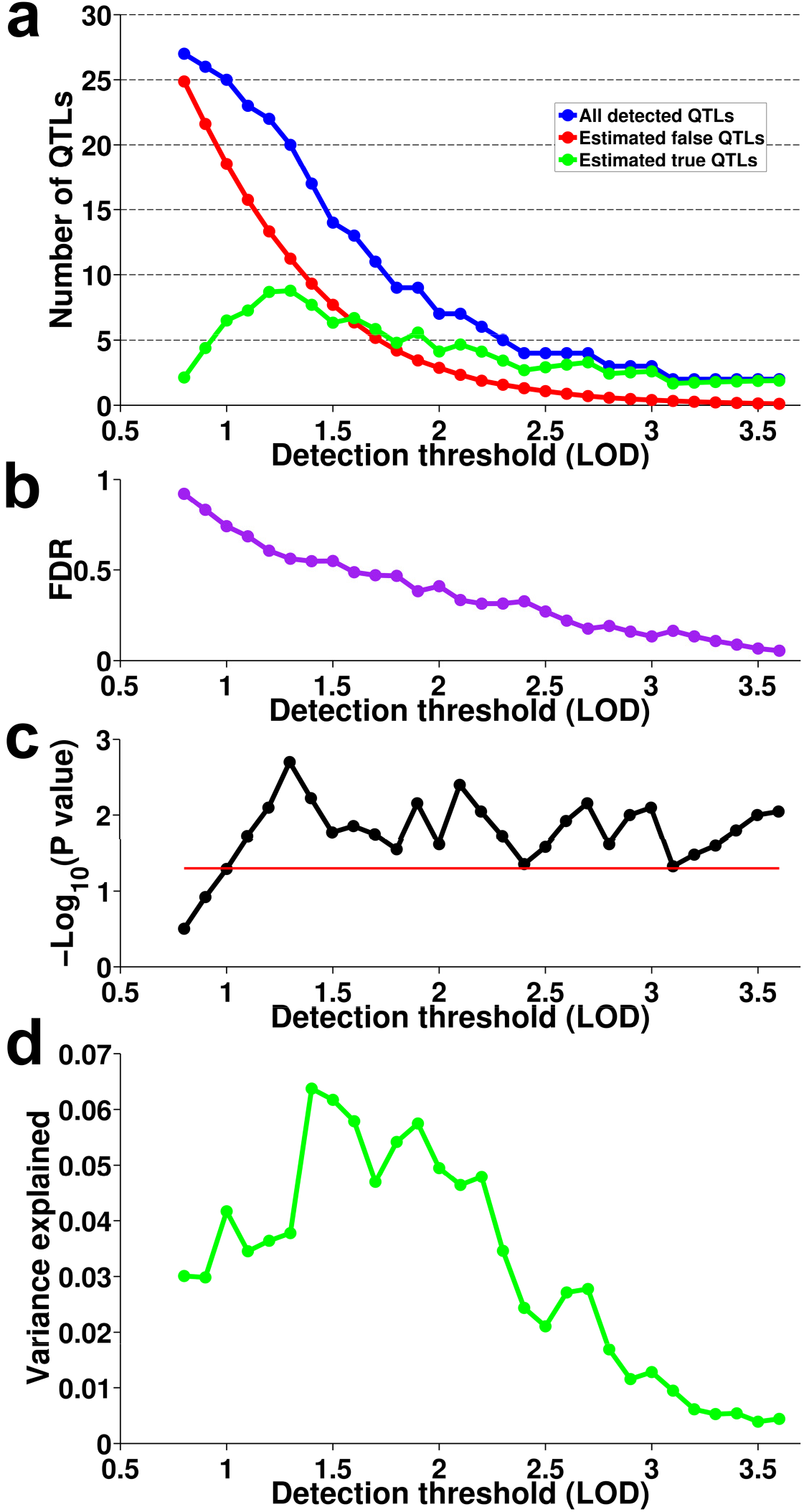
QTL mapping. **(a)** For each LOD detection threshold, the number of detected QTLs is plotted in blue, the expected number of false-positive QTLs based on permutation analysis is plotted in red, and the difference between these two numbers is plotted in green. **(b)** Permutation-based false discovery rate at each detection threshold **(c)** Statistical significance of the number of detected QTLs at each threshold, shown as -Log10(permutation-based P-value). Red line shows P=0.05. **(d)** Variance explained by QTLs in a cross-validation framework.

### Variance explained by QTLs

To estimate how much phenotypic variance can be explained by the detected QTLs, and to test whether the detected loci tag common SNPs identified by previous height GWAS ^47^, we conducted variance partitioning in a cross validation framework. We generated 100 training sets that each randomly sampled 2/3 of the families. In each set, we mapped QTLs as described above. The results were similar to those obtained with the full dataset (**Fig. S6**), although on average, fewer QTLs were in the training sets as a consequence of the smaller sample size. We then used the 1/3 of the families held out of each training set as a test set, and simultaneously estimated the contributions of three different sources of genetic variance to height variance in a variance components model (implemented in the software package GCTA ^48^). Specifically, the model included variance component terms for the detected QTLs, the common SNPs associated with height by GWAS ^47^, and the overall genomic relatedness among individuals; the latter controls for pedigree structure. We also considered a null model in which the QTLs have no effect on height but simply capture some genomic relatedness; we estimated the expected variance explained by the QTLs in this model by running GCTA as above but with random segments of the genome that match the QTLs in length (**Methods M12**).

As expected, total variance explained by the QTLs increased as the detection threshold was lowered and a larger number of QTLs were included in the model, but the variance explained over and above the null model initially also increased (**Fig. 2d**). For example, at LOD≥3, we detected an average of 0.74 estimated true QTLs per training set (1 QTL at FDR=26%), and these explained an average of 1.3% of the phenotypic variance above the null model in the test sets. At a lower detection threshold of LOD=1.9, we estimated an average of 3.7 true QTLs (7 QTLs at FDR = 47.5%), and these explained 5.8% of the variance above the null model. For low detection thresholds (LOD≤1.3), the variance explained above the null model decreased, presumably because too many false discoveries were included.

To test whether QTLs explained variance in height by tagging previously discovered height-associated common variants, we ran the variance components model with and without including the GWAS SNPs. The variance explained by the QTLs in both models was similar (0.1% difference, t-test P = 0.72, **Fig. S7-S8**). This result suggests that the QTLs we identified are novel and are not simply tagging common SNPs that were previously identified by GWAS.

### Specific chromosomes contribute disproportionally to height and in accordance with the QTL signals

To assess how much phenotypic variance can be explained by entire chromosomes, we estimated a genomic relatedness matrix (GRM) from all the SNPs on each autosome, and let the 22 GRMs compete in a variance components model to explain phenotypic variance (**Fig. 3**). The sum of variance explained by all chromosomes was 92%. We found no correlation between variance explained and chromosome length (Pearson r = -0.08, P = 0.71), although we cannot rule out that such a correlation exists, as the standard errors in the estimation of single chromosome contributions are large. In contrast, the variance explained by chromosomes is highly correlated with the top LOD score on each chromosome (r = 0.7, P = 2.9×10^-4^). This correlation arises in part from the results for chromosome 14, which explains the most variance in the variance components model (25%±7.8%) and has the single most significant QTL. To test whether the correlation is driven solely by chromosome 14, we omitted it from the analysis. The correlation fell but remained significant (r = 0.53, P = 0.015).

These results suggest that at least in our sample, variance explained by some of the chromosomes captures contributions of small regions with large effects rather than solely infinitesimal contributions distributed throughout the entire chromosomes. To investigate this further, we simulated 100 sets of phenotypes from an infinitesimal model, in which normally distributed small effect sizes were randomly assigned to all SNPs in the genome while maintaining the overall heritability. For each simulated data set, we calculated the distribution of variance explained by entire chromosomes. The correlation between variance explained and chromosome length had on average of r=0.19, higher than the r = -0.08 observed for the real data, although the difference was not statistically significant (P = 0.09, **Fig. S9b**). The observed variance explained by chromosome 14 was significantly higher than expected from the infinitesimal model. Only 5/2200 chromosomes in the simulated data sets explained as much as 25% of the variance (Bonferroni corrected P = 0.05, **Fig. 3c**), and all of these five observations were for chromosomes that are longer than chromosome 14.

**Fig. 3.**
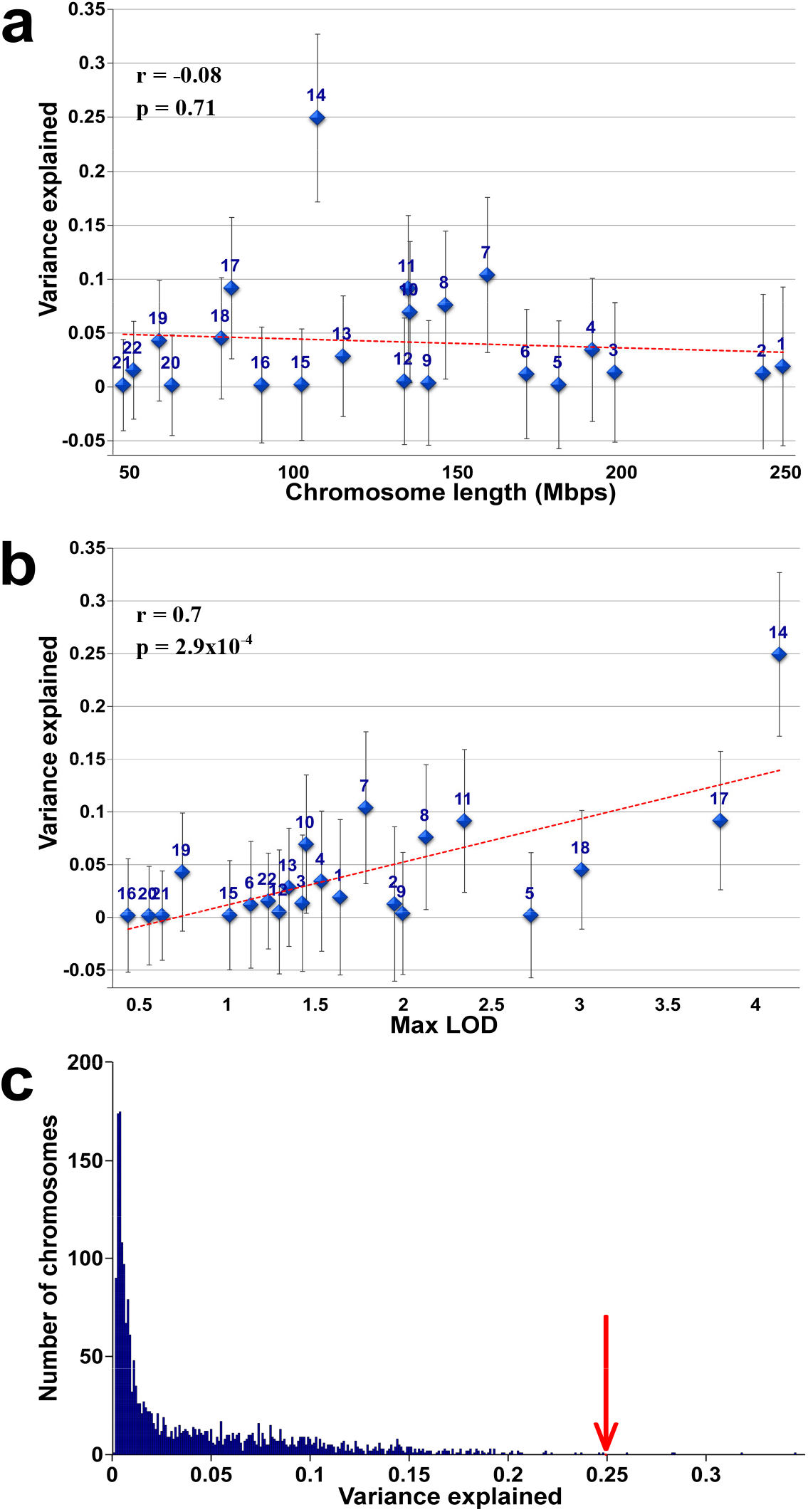
Height variance explained by chromosomes. **(a)** Height variance explained by each chromosome ± SE is plotted against chromosome length. Red line shows the linear fit. **(b)** Height variance explained by each chromosome ± SE is plotted against the maximum LOD score on that chromosome. **(c)** Histogram of variance explained by each of 2200 chromosomes simulated under an infinitesimal model. Red arrow shows variance explained by chromosome 14 in the real data.

## Discussion

Here, we studied height in a unique cohort of very large nuclear families from a founder population. This strategy was designed to increase the effective allele frequency of some variants that are otherwise rare, thereby also increasing our power to detect their effects on height. This approach enabled us to detect significant QTLs for height in a study with modest sample size. We also used FDR analysis to identify a larger number of QTLs that were highly significant as a set, despite few of the QTLs achieving significance individually. Using a variance components model in a cross-validation framework, we showed that these QTLs explained 6% of the variance in height in our sample, and that they were not tagging common variants previously identified as associated with height by GWAS. The actual fraction of variance explained by the QTLs is likely higher because of the conservative nature of the estimation procedure. Further, we showed that the variance contributed by chromosome 14, and possibly by some of the other chromosomes, arises at least in part from small regions with large effects (which correspond to the detected QTLs), rather than solely from infinitesimal contributions distributed throughout the entire chromosomes.

Taken together, these results suggest that variation in height in our sample arises from a combination of a small number of QTLs with large effects and a large number of common variants with small effects. Because the detected QTLs are not tagging previously identified common variants, they likely arise from variants that are elevated in frequency in the AJ population. Although we have not identified the specific variants underlying the QTLs, we speculate that candidate variants can be identified by sequencing the parents of the pedigrees and searching for variants that are rare in other populations, common in AJ, and follow segregation patterns consistent with the QTL signals. The approach described in this paper, coupled with recruitment of additional large families (which are abundant in the Jewish population ^49,50^), may provide further insights into the genetic basis of height and the role of population-specific vs. cosmopolitan variants, and may serve as a complement to GWAS for genetic investigations of other complex traits.

## Acknowledgments

We thank Kruglyak laboratory members for helpful discussion, Eran Segal for lab space and equipment, and the Puah Institute for help in the recruitment of participants. Funding was provided by the Howard Hughes Medical Institute (L.K. and D.Ze), and the James S. McDonnell Centennial Fellowship in Human Genetics (L.K.).

